# A geography aware reconciliation method to investigate diversification patterns in host/parasite interactions

**DOI:** 10.1101/166215

**Authors:** V. Berry, F. Chevenet, J-P. Doyon, E. Jousselin

## Abstract

Cospeciation studies aim at investigating whether hosts and symbionts speciate simultaneously or whether the associations diversify through host shifts. This problem is often tackled through reconciliation analyses that map the symbiont phylogeny onto the host phylogeny by mixing different types of diversification events. These reconciliations can be difficult to interpret and not always biologically realistic. Researchers have underlined that the biogeographic histories of both hosts and symbionts influence the probability of cospeciation and host switches, but up to now no reconciliation software integrates geographic data. We present a new functionality in the *Mowgli* software that bridges this gap. The user can provide geographic information on both the host and symbiont extant and ancestral taxa. Constraints in the reconciliation algorithm have been implemented to generate biologically realistic codiversification scenarios. We apply our method to the fig/fig wasp association and infer diversification scenarios that differ from reconciliations ignoring geographic information. In addition, we updated the reconciliation viewer *SylvX* in order to visualize ancestral character states on the phylogenetic trees and highlight zones that are geographically inconsistent in reconciliations computed without geographic constraints. We suggest that the comparison of reconciliations obtained with and without constraints can help solving ambiguities in the biogeographic histories of the partners. With the development of robust methods in historical biogeography and the advent of next-generation sequencing that leads to better-resolved trees, a geography aware reconciliation method represents a substantial advance that is likely to be useful to researchers studying the evolution of biotic interactions and biogeography.

## 1) INTRODUCTION

Biotic interactions play a prominent role in species diversification. Interactions that result into long-term associations persisting over evolutionary time scales can sometimes lead to cospeciation, i.e. the concomitant occurrence of speciation in lineages that are ecologically associated (Brooks 1981; Page 1990, 1991). The idea that such a pattern can occur first stemmed from parasitological studies suggesting that parasite classifications reflect the phylogenetic relationships of their hosts (Fahrenholz 1913). Hafner & coll. (Hafner & Nadler 1988; Hafner *et al.* 1994) were the first authors to thoroughly test this assertion. They used the association between pocket gophers and their lice as a model system and provided a clear demonstration that the phylogenies of the two interacting lineages were parallel. This study spurred further research on cospeciation. The developments of specific methods that aimed at testing the congruence of the phylogenetic histories of interacting organisms have since played an important role in the study of cospeciation. It is indeed these methods that moved cospeciation studies beyond visual comparisons of phylogenetic trees and ad-hoc narratives for these visualizations. It soon became apparent that the study of the concordance between phylogenetic trees could be applied to reconciling gene trees and species trees (Page & Charleston 1997; Page & Charleston 1998) which further enhanced the interest of evolutionary biologists for methodological developments in this field.

Reviews on cospeciation methods (Brooks *et al.* 2004; de Vienne *et al.* 2013; Doyon *et al.* 2011; Johnson & Clayton 2004; Martínez-Aquino 2016; Paterson & Banks 2001; Stevens 2004) all emphasize the diversity and the complexity of the scenarios that must be explored when testing for the congruence of speciation events in two interacting lineages. To compare host and parasite phylogenies, *Brooks & coll.* (Brooks 1981; Brooks & McLennan 1991) first developed a parsimony method (the Brooks Parsimony Analysis, BPA). In this method the associations between hosts and their parasites are transformed into a matrix of host characters and the parsimony tree reconstructed from such a matrix is then compared to the host phylogeny. A decade later, Page and collaborators developed a fundamentally different method, called “tree reconciliation”, a term first coined in the work of Goodman *et al.* (1979) that compared gene and species trees. This method attempts to reconcile the phylogenetic history of the parasite with that of their hosts: the parasite phylogeny is “mapped” onto the host phylogeny (i.e. each node in the parasite tree is assigned to a node or a branch in the host phylogeny). In such a map, the diversification events of the parasites are linked to their host phylogenetic history and four types of events are considered: cospeciation events, host switches, sorting events and duplication events (Page 1994a; Page 1994b) (see material and method for a description of each event). When graphically displayed, reconciliation maps greatly ease our understanding of the evolution of biotic interactions.

Algorithms to optimize reconciliations are numerous. One of the first reconciliation software, *TreeMap* 2, uses an algorithm called “Jungles” (Charleston 1998) where each event is assigned a cost: the chosen reconciliations are the ones that have minimum costs. However it generates in the process an exponential number of scenarios. Recent methods have proposed algorithms that are more efficient and can also just search for an optimal reconciliation: e.g. *Tarzan*, (Merkle & Middendorf, 2005); *Jane*, (Conow *et al.* 2010); *Core-PA* (Merkle *et al.*, 2010), *Mowgli*, (Doyon *et al.* 2010), *COALA* (Baudet *et al.* 2015), ecce*TERA* (Jacox *et al.* 2016), *Notung* (Stolzer *et al.* 2012), EUCALYPT (Donati *et al.* 2015) and ILPEACE (van Iersel *et al.* 2014). Recently, the *RASCAL* software proposed to infer suboptimal scenarios to reduce computing times (Drinkwater & Charleston 2016). Cospeciation is witnessed on a reconciliation map whenever a speciation node in the parasite phylogeny is mapped onto a speciation on the host phylogeny. Another requirement for demonstrating that two interacting lineages have cospeciated is to provide evidence of the temporal congruence of the cospeciation event in the host and parasite phylogenies (Page 1991). Though reconciliation algorithms do not strictly enforce the simultaneity of cospeciation events, they can enforce time consistency in the sequence of evolutionary events, meaning that the parasite cannot switch back in time onto a host that no longer exists (*i.e.* transfers cannot occur towards a node in the host phylogeny that has already split into child species at the time of the transfer event) (Merkle & Middendorf 2005; Nøjgaard *et al.* 2017). This constraint is explicit in *Mowgli* (Doyon *et al.* 2011; Doyon *et al.* 2010), *ecceTera* (Jacox *et al.* 2016) and *RASCAL* (Drinkwater & Charleston 2016). Hence, reconciliation methods have greatly improved in the last decade; algorithms are now efficient and some have solved the time consistency issue that affected some of the first methodological developments in the field. However, interpreting the scenarios that emerge from these inferences remains a difficult task. It is generally challenging to identify biologically realistic reconciliations. Much remains to be done to improve these inferences and translate them into evolutionary scenarios that give insights into the biological factors that govern the evolution of interspecific associations.

Some key information that could significantly improve our inferences but are overlooked in codiversification methods are the geographic locations of extant and ancestral nodes. Indeed, the biogeographic histories of interacting lineages necessarily constrain their common part of evolutionary history (Martinez-Aquino *et al.* 2014; Nieberding *et al.* 2010). Obviously, a cospeciation event can only happen between taxa that co-occur in the same area. The geographic context of both hosts and parasites also influences host switch events. In biotic interactions where the parasites can undergo long dispersal events, transfers can happen between allopatric hosts (i.e. hosts that do not live in the same geographic area). However, they are only possible if the geographic locations of the “sending host” (the host from which the switch is initiated) and the “receiving host” coincide with a dispersal event along the corresponding branch in the parasite phylogeny. Therefore, a more accurate mapping of cospeciation and host switch events can be obtained if the geographic locations of both hosts and parasites are known prior to conducting the reconciliation.

Methods for inferring historical biogeography from phylogenetic reconstructions have greatly improved in the last two decades. Early developments in historical biogeography aimed at reconstructing “area cladograms” that reflected the history of connections between areas of endemism for the group of organisms under study and used analytical tools that were very similar to the tools developed for the study of cospeciation using parsimony as the optimization criterion (*e.g.*, BPA, see Morrone, 2009 for a review on cladistic biogeography and its methodological developments). More recent probabilistic methods in the field of historical biogeography aim at reconstructing ancestral geographic range of focal lineages from current species distribution and a dated phylogenetic tree. They model the evolution of geographic areas on a phylogenetic tree using Maximum Likelihood optimization or Bayesian inference and incorporate divergence times into the inference process: the longer the phylogenetic branch, the higher the probability of geographic range shifts and the larger the uncertainty in the ancestral range estimates. Geographic areas can be treated as simple categorical characters that are reconstructed on the tree using for instance a stochastic Markov model of evolution. More biologically realistic and widely applied methods in historical biogeography, such as DEC (*Dispersal, Extinction, Cladogenesis*) (Ree *et al.* 2005; Ree & Smith 2008), model range evolution using different parameters for each biogeographic process (dispersal, range expansion or extinction). In addition to modelling these key processes in range evolution, the main innovation of DEC consists in incorporating a time-dependent transition matrix that defines the movements between geographic areas, at different time intervals in order to reflect how dispersal opportunities changed through time (*e.g.* changes in the continents configuration for instance) (see Ree & Sanmartin 2009; Ronquist *et al.* 2011, for reviews on parametric biogeography). Fossil distribution and information on the climatic preferences of ancestral lineages can also be incorporated as constraints to improve biogeographic inferences (Meseguer *et al.* 2015). Several conceptual and computational improvements have been implemented since the initial version of DEC (DEC + J, Matzke 2014; DECX, Beeravolu Reddy & Condamine 2016). Different biogeographic models have also been proposed (GeoSSE, Goldberg *et al.* 2011; BayArea Landis *et al.* 2013). As a result, robust biogeographic scenarios are now available for numerous lineages. Ancestral areas inferred by these methods can then serve as input for reconciliation analyses. In this paper we build on these advances to provide a geography-aware reconciliation method, pushing further the realism of scenarios proposed by such methods.

We first describe the constraints we enforce to ensure geographic consistency in reconciliations and how they were implemented in the *Mowgli* reconciliation software (Doyon *et al.* 2010). We also updated the *SylvX* reconciliation viewer (Chevenet *et al.* 2016), in order to integrate and visualize annotations (e.g. geographic areas) at ancestral nodes for the host and parasite phylogenies and highlight inconsistent zones in the reconciliation. We then test these new developments on a mock dataset and on a ‘textbook’ example of cospeciation, namely the interaction between figs (*Ficus*) and their pollinating fig wasps (Cruaud *et al.* 2012; Rønsted *et al.* 2005; Wiebes 1979).

## 2) METHODS

### Extending *Mowgli* to account for geographic information

In this section we first recall the reconciliation model followed by *Mowgli* (Doyon *et al.* 2010).

Only rooted parasite and host trees are considered; their leaf nodes (tips) are each labelled by a taxon name. The host tree is dated, meaning that either each branch length represents an amount of time (the tree is thus ultrametric) or that the age of each internal node is provided (e.g. in million years). Internal nodes usually have two descendants, but an internal node can also have a single child also when the evolution of an ancestral lineage living a relatively long period of time is decomposed into a set of consecutive time periods called *slices* (see Fig. 3 of *Mowgli* Manual). This slicing of branches is a transparent artefact that allows reconciliation methods to achieve fast computing times while still ensuring time consistency of host switches (see Doyon et al. 2010; Jacox *et al.* 2016; Libeskind-Hadas & Charleston 2009).

Let *P* and *H* denote respectively a parasite and a host tree, *x* and *x_p_* are nodes (or extant species, *i.e.*, leaves) of *H* and *u* and *u_p_* are nodes (or extant species) of *P*. Reconciliation algorithms usually consider each current and ancestral host to be associated with one or several specific parasites at any time (*e.g.* in *Mowgli*, *TreeMap*, *Jane*). However, the identity of the host can vary over time, *e.g.* after a *host switch*. This evolutionary event is one of the four types of *events* considered in cospeciation studies:

- a *host switch*, also known as a transfer (T event), occurs when a parasite lineage from a source host is transferred to a destination host. The transfer of the parasite must be time consistent, that is the “sending” branch (*x_p,_x*) and the “receiving” branch (*x’_p,_x’*), where the host switch is mapped, must belong to the same time slice;
- a *cospeciation* (S event) happens when the speciation of a parasite shortly follows or coincides with the speciation of its host. This is considered by *Mowgli* as a joint speciation of both parasite and its host;
- a *within host speciation* also known as duplication (D event), models a speciation of a parasite *u* of *P*, where both descendant species continue to live on the host that *u* lived on. This is represented by *u* evolving along a (*x_p,_x*) branch of H and then splitting into two new lineages in (*x_p,_x*);
- a parasite *loss* (L event) occurs when a parasite lineage goes extinct while its host persists.

An illustration of these events can be found the *Mowgli* Manual.

*Mowgli* also sometimes considers combinations of events in order to speed up computations. A SL event occurs shortly after a cospeciation (S): one of the parasite child lineages is quickly lost (L) in the host phylogeny on a child lineage of the involved speciation node. A TL event occurs when a parasite *u* evolving on a branch (*x_p,_x*) is lost (L) on this branch shortly after having switched (T) to another host (*x’_p,_x’*).

As explained above, accounting for geographic information can lead to more realistic diversification scenarios. We first integrate such information by assigning a set of *areas* to each node of *P* and *H*. For an extant taxon this means that a population of the corresponding species is reported to live in *each* of the assigned areas. In contrast, when an internal taxon is assigned to one or several areas, this means that populations of this now extinct taxon are inferred to have lived back in time in one or several of these geographical zones.

In order to compute biogeographically meaningful reconciliations between the *P* and *H* trees, specific constraints have to be implemented in reconciliation algorithms. We detail below how we model these constraints in the context of the four D/T/L/S events or combinations thereof. First, note that areas of a node and its parent in the host or parasite tree can be different, due to dispersal and vicariance events. During the reconciliation process, the time period represented by a branch between nodes *x_p_* and *x* of the host tree is considered to be assigned the union of the areas of *x_p_* and *x*. If a species changes area along the branch from one area assigned to *x_p_* to a different one assigned to *x*, we do not know exactly when it happened, so we consider that at any time between *x_p_* and *x*, part of the population of the evolving species can live in any area proposed for *x_p_* or *x.*

Considering nodes of the trees, we denote by *area*(*x*) the set of geographic areas where an extant species *x* is observed (at the tip of a tree). Areas proposed for an internal node *x*, that is for an extinct species, are also denoted *area*(*x*). However, as indicated above, the meaning is somewhat different as *area*(*x*) represents in this case the set of areas where *x* could have lived. Because of the incertitude in the historical biogeography inferences, we do not enforce that *x* lived in each of these areas. Similarly, considering branches (*x_p_,x*) of the *H* tree, *area*(*x_p_,x*) denotes the set of areas where the species might have lived during this period: this is the union of *areas*(*x_p_*) and *area*(*x*). Note that each area in which exactly one of the two species *x* and *x_p_* is present corresponds to a migration or extinction event that has occurred along this branch. In addition, only (*x_p_,x*) branches being one slice higher are considered for *H*, as *Mowgli* operates on this level of detail.

We now detail which geographic constraints apply so that the reconciliation between a parasite tree *P* and a host tree *H* is geographically consistent. Recall that a reconciliation is a mapping of *P*’s nodes and branches onto those of *H*.

- An extant parasite *u* can be mapped onto an extant host *x*, only if *area*(*u*) ⊆ *area*(*x*) (Fig. 1 A). If this constraint is not fulfilled then *Mowgli* cannot compute a reconciliation.
- We allow the mapping of an ancestral parasite *u* at a speciation node *x* in the host tree, only if *area*(*u*) and *area*(*x*) have a non-empty intersection, *i.e.* when there is at least one area where the parasite and the host were able to meet (Fig. 1 B).
- A parasite node *u* can be mapped into a branch (*x_p_,x*) of *H* due to a duplication or host switch event (Fig. 1 C), and in those cases, we also require that *area*(*u*) ∩ *area*(*x_p_,x*) ≠ ∅. Note, that this constraint does not prevent parasite dispersal events during host switches.
- If a branch (*u_p_,u*) of the parasite tree is mapped for all or part of it onto a host branch (*x_p_,x*) (Fig. 1 D), then we also require that *area*(*u_p_,u*) ∩ *area*(*x_p_,x*) ≠ ∅.
- Last, if a branch (*u_p_,u*) of the parasite is going through a node *x* of the host tree (which happens when the host speciates into two descendant hosts but the parasite sticks to only one of them – an SL event), then the area(*x*) and area(*u_p_,u*) must have common elements (Fig. 1 E).

**Fig. 1:**
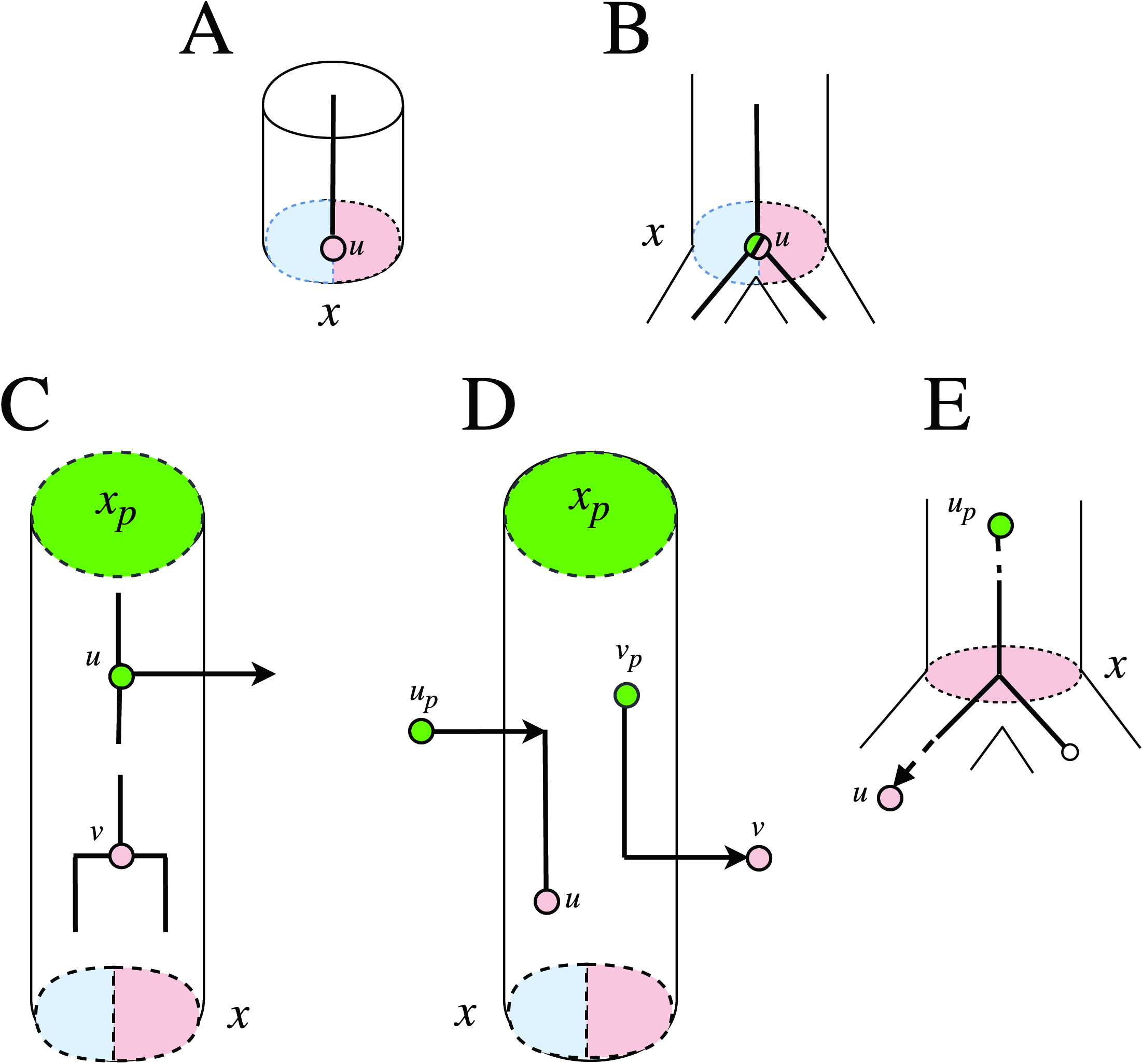
Description of how geographical constraints are handled by the *Mowgli* software. Plain lines and nodes represent branches and nodes of the parasite tree, while cylindres and dashed ellipses represent branches and nodes of the host tree. The colours of a node correspond to geographical areas, these areas are observed (hence enforced) for extant taxa but inferred for ancestral nodes. A) The parasite tip *u* can be mapped to a tip *x* of the host tree if the areas of the host contain all areas of the parasite. B) *Mowgli* accepts that a parasite node *u* cospeciates with a host at a node *x*, if the two nodes share an area. Here, it was inferred that the ancestral parasite *u* lived in green and/or red areas, but cospeciated with a host *x* that lived in blue and/or red areas, we conclude that *u* lived only in the red area at this time and that *x* lived at least in the red area. C) To map an ancestral parasite inside a branch (*x_p_,x*) – to represent the source of a host switch (upper part of the figure), or a duplication of the parasite (lower part) — *Mowgli* requires that the parasite has potentially lived in an area of *x_p_* or *x*. In this example, *u* shares an area with *x_p_* and *v* shares an area with *x*. D) To map a branch (*u_p_,u*) of the parasite tree inside a branch (*x_p_,x*) of the host tree, *Mowgli* requires that the parasite mapped on the host branch (*u* in the left part of the figure showing the destination of a switch and *v_p_* in the right part showing the departure due to a switch) has potentially lived in any area assigned to node *x_p_* or to node *x*. This is the case here for node *u* that was indicated as having lived in the red area (also assigned to *x*) and for *v_p_* assigned to the green area, also proposed for *x_p_*. Note that mapping *v_p_* into (*x_p_,x*) would have also been correct if *v_p_* had been assigned to the red area, indicating that it changed from the green to the red area, together with its host, before switching to another host. E) When a parasite lineage (*u_p_,u*) living on an ancestral host remains with one descending child of this host after its speciation at node *x*: *Mowgli* requires that the area at which the host speciation occurred is also found among the areas inferred for *u_p_* or *u.* The mapping in this example indicates that the parasite changed area with its host, before the host speciation event.

Note that when part of the reconciliation mapping traverses an artificial node *x* in H, then no particular constraint applies locally: the possibility of such a scenario is directed by constraints ensured with respect to the branch (*x_p_,x*) of *H* to which *x* belongs.

When respecting the above constraints, *Mowgli* will propose a scenario that is geographically consistent. This scenario can have a higher cost than those obtained when not accounting for geographic information. This simply results from the fact that the search space contains geographically inconsistent scenarios that are possibly less costly. *Mowgli*’s extension described above, allows choosing the less costly scenario among those that are geographically consistent.

### *SylvX’s* new functionalities

We extended the *SylvX* editor in order to visualize current and ancestral geographic areas of hosts and symbionts. Pie charts can be used to display alternative areas for each node of the tree and/or the reconciliation. Area colour sets can be dynamically updated and tuned using the Hue, Saturation and Value scales. Thresholds are available to simplify views. *SylvX* also contains a new tool in the *Annotation* panel to highlight reconciliation parts that do not respect geographical constraints (when such constraints have not been enforced when computing the reconciliation). This is done by loading an annotation file generated by *Mowgli* (constraintsPBM.csv).

#### Implementation

*Mowgli* takes as input a “host tree” and a “parasite tree” stored in files in a Newick format. A list of nodes with their geographic areas (or other annotations) can be given in the same files. Biogeographic inferences typically generate probability or likelihood values for each character state (area) at each node. *Mowgli* can accept a single area or a set of areas at each node. To run *Mowgli* and obtain a reconciliation respecting geographical constraints, the–a flag must be added in the command launching the program. Adding the–y flag instead computes a reconciliation independently of the indicated constraints but pinpoints the places where the mapping violates these constraints (in `mapping.mpr` and `constraintPBMs.csv` files, see the provided manual for details). This allows users to identify inconsistencies between the most parsimonious reconciliations and the hosts and parasites respective biogeographic histories.

*SylvX* takes a host tree in Newick format with node *id* numbers and a reconciliation (with symbiont tree node *id*.). The host tree (`outputSpeciesTree.mpr`) and reconciliation obtained with *Mowgli* can be directly imported into *SylvX.* The latter also supports input files from other reconciliation software, *e.g.*, ecceTERA (Jacox *et al.* 2016). Annotation files for the host and parasite phylogenies giving node information can be imported in a CSV format. As many annotations as needed can be added in the annotation files and it is up to the user to choose which ones to plot onto the species tree and the reconciliation map through *SylvX*’s interface.

In order to seamlessly pass a user annotation file in csv format into both *Mowgli* and *SylvX*, we provide a *Perl* script that can be run through the command line in order to: 1) obtain tree node identifiers that will be used by both programs and 2) merge input trees and corresponding annotations files into *Mowgli’s* input format. Files can be generated so that a single (most likely) ancestral range can be specified or alternative geographic areas can be assigned to all nodes (see Supplementary Material 1 for a description of the full procedure to generate files, set a threshold value above which to keep alternative areas and perform a complete analysis).

## 3) WORKED EXAMPLE

### Datasets

To demonstrate the method and its utility, we tested it on two datasets. We first created a mock dataset: two phylogenetic trees with nine tips for a hypothetical host/parasite interaction in which extant and ancient geographic areas for each lineage are informed. The dataset was generated by hand so that: 1) present-day geographic areas of associated taxa are consistent (*i.e.*, hosts and associated parasites live in the same area); 2) the two phylogenies are not perfectly parallel but show some cospeciation events; 3) some geographic locations at nodes that we would like to cospeciate do not coincide in the parasite and host phylogenies. We ran *Mowgli* on this dataset successively with and without enforcing geographic constraints using in both cases the default parameters (cost 0 for a cospeciation, 1 for a loss and 1 for a host switch, 1 for duplication, not enforcing the root of the parasite tree to be mapped on the root of the species tree). In order to measure the impact of cost settings on the reconciliation scenarios, we ran this dataset using alternative costs for host switches and losses.

As a second dataset, we used a subset of the data from the latest phylogenetic investigation of figs (*Ficus*) and their pollinating wasps (Cruaud *et al.* 2012). For both partners of the association, biogeographic scenarios were available for phylogenies of 200 taxa. From the complete phylogenetic trees (available in http://datadryad.org, doi: 10.5061/dryad.hr620), we derived two trees of 23 taxa each, that included a couple of representative species for each *Ficus* main taxonomic subdivision. We excluded one of the fig subgenera (*Pharmacosycea*) and its associated pollinators (*Tetrapus* spp.) whose phylogenetic positions are still debated. We have not tested our method on the total dataset presented in Cruaud *et al.* (2012) as some uncertainties remain concerning the root of the phylogenetic trees, which could lead to spurious interpretations. The most likely ancestral geographic areas of each node were directly derived from the biogeographic reconstructions of Cruaud *et al.* 2012, obtained with Maximum Likelihood Optimization in Mesquite (Maddison & Maddison 2006). We ran *Mowgli* on this dataset with and without enforcing geographic constraints (using default event costs and not enforcing the root of the parasite tree to be mapped on the root of the species tree), and explored how these reconstructions shed light on the biogeographic history of the association. In order, to investigate how incertitude on ancestral geographic ranges impacts the reconciliation, we ran the reconciliation on the dataset including alternative ancestral areas for both *Ficus* and their associated pollinators. For each node of the pollinator and the *Ficus* phylogenies, the geographic areas which proportional likelihood was above 0.15 were kept and assigned to their respective nodes.

### Results

Figure 2 represents the reconstruction obtained on the mock dataset. When not taking geographic constraints into account (Fig. 2A), a cospeciation event at a node where the two associates do not live in the same area was retrieved (node S1 of the host tree in Fig. 2A). The transfer T1 preceding this cospeciation event is also geographically impossible as it suggests a dispersal (the donor host lives in Asia or Africa, and the receiving host lives in America) while the parasite actually stays in Africa. The scenario obtained when enforcing geographic constraints is more costly (Fig. 2B): it entails one more transfer and consequently one less cospeciation event but is biologically more realistic. When using different cost vectors (i.e. using a cost of 3 for losses), the reconciliation where geographic constraints are not taken into account includes additional transfers to avoid losses (Fig. S2A); those are all geographically inconsistent. The reconciliation with geographic constraints also changes (Fig. S2C) and necessitates 5 transfers to ensure geographic consistency of the diversification events in hosts and parasites without inferring any parasite losses. When we increased the cost of transfers (cost T=10, Fig. S2B), the reconciliation without geographic constraints infers several early duplications and losses in order to avoid a costly transfer. On the other hand the results of the reconciliation under constraints (Fig. S2D) did not change comparatively to the one obtained with default cost settings. Hence, in this particular case, adding biological constraints into the reconciliation process stabilizes the reconciliation and makes it less dependent on cost settings.

**Fig. 2:**
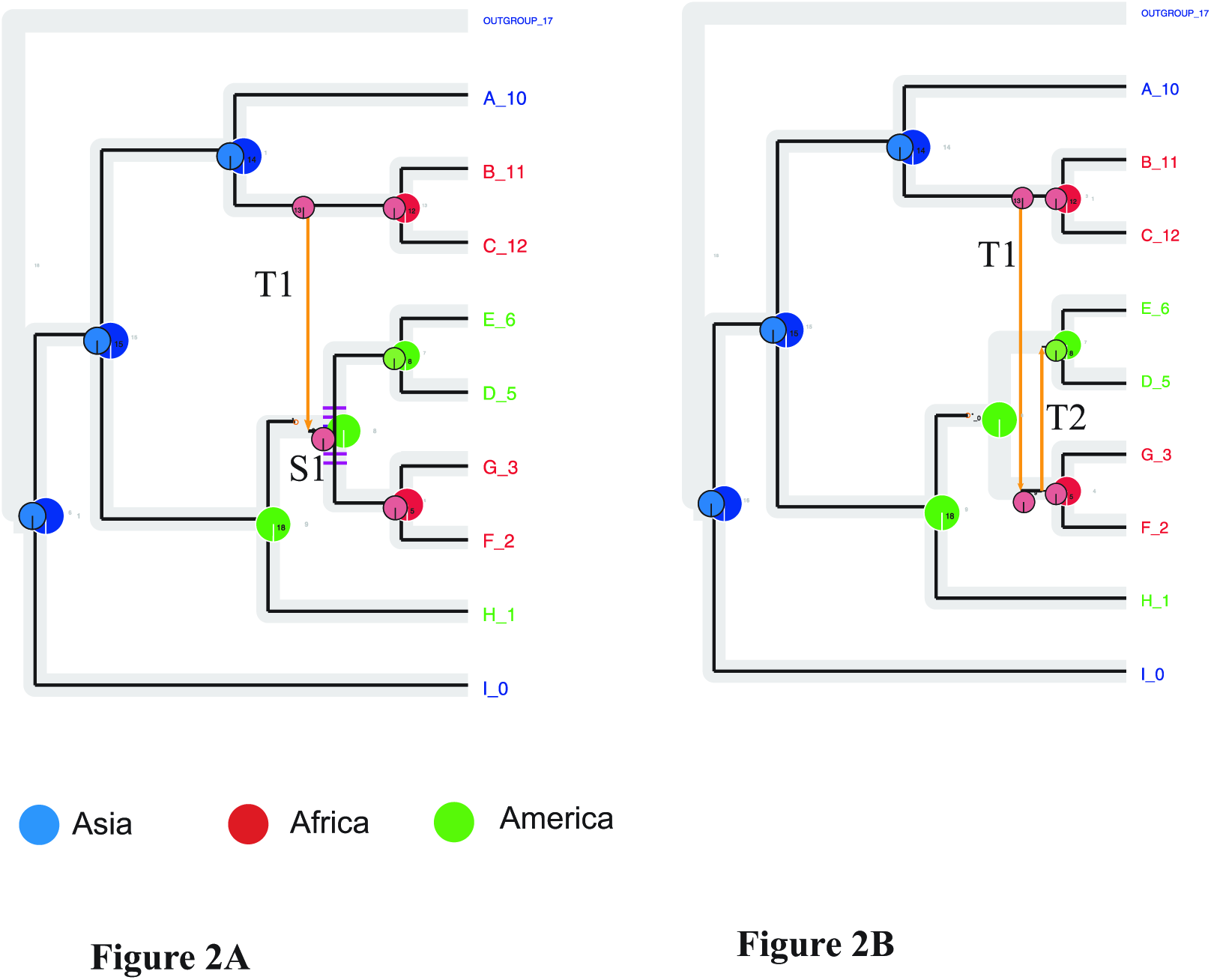
Results of the reconciliations obtained on a simulated data set with default cost settings: A) without enforcing geographic constraints (7 coSpeciation events, 1 Transfer, 1 Loss), purple dashed branches indicate parts of the reconciliation where geographic constraints are not fulfilledB) when enforcing geographic constraints (6 S, 2 T, 1 L). In both cases, the big pie charts correspond to the host ancestral geographic areas while small pie charts correspond to the symbiont ancestral geographic areas. The leaves of the species tree are also coloured according to the current geographic distribution of the associates. Annotations files given for the host tree and the symbiont tree specified a single most likely area at each node.

On the fig/fig wasp dataset (Fig. 3), not accounting for geographical constraints leads to geographic inconsistency in one node (cospeciation S1 in the host tree of Figure 3A). The transfer that precedes it (T5) is impossible and the mapping of the fig wasp phylogeny onto the fig phylogeny from node S1 to event T6 is geographically inconsistent ((Fig. 3A). Enforcing geographic constraints when a single (most likely) area is specified for each node generates a reconciliation scenario that is more costly (Fig. 3B, one more transfer is necessary to reconcile the two phylogenies) but coherent with the figs and the fig wasp biogeographic histories. This scenario suggests that the fig wasps independently colonized figs in the Neotropics and in the Afrotropics through two distinct host switches from Asia rather than accompanied the speciation of their hosts, as was suggested by Figure 3A (and node 29 of Fig. S12 in Cruaud *et al.* 2012). The annotation of ancestral geographic areas on the reconciliation map also shows that host switches occurred both in “sympatric” settings (within the same geographic areas as broadly defined in our dataset) and allopatric settings (i.e. host switches occur between two geographically distant hosts). Overall, four switches out of seven occurred in sympatry (T1, T2, T4, T5) while the remaining three switches (T3, T6, T7) correspond to long distance dispersal events (Fig. 3B). Adding incertitude in ancestral geographic range, generates a reconciliation that matches the one obtained without constraint (Fig. 3C), as geographic areas of node S1 of the host figs now includes Asia among its potential geographic areas. This matches the ancestral geographic area of the inferred associated fig wasps. In that scenario a single host switch is associated with a long dispersal event of the fig wasps (T3: from Asia to Australasia), all other host switches occur in sympatric settings (within Asia) and fig wasp geographic range evolution merely mirrors the one of their hosts.

**Fig. 3:**
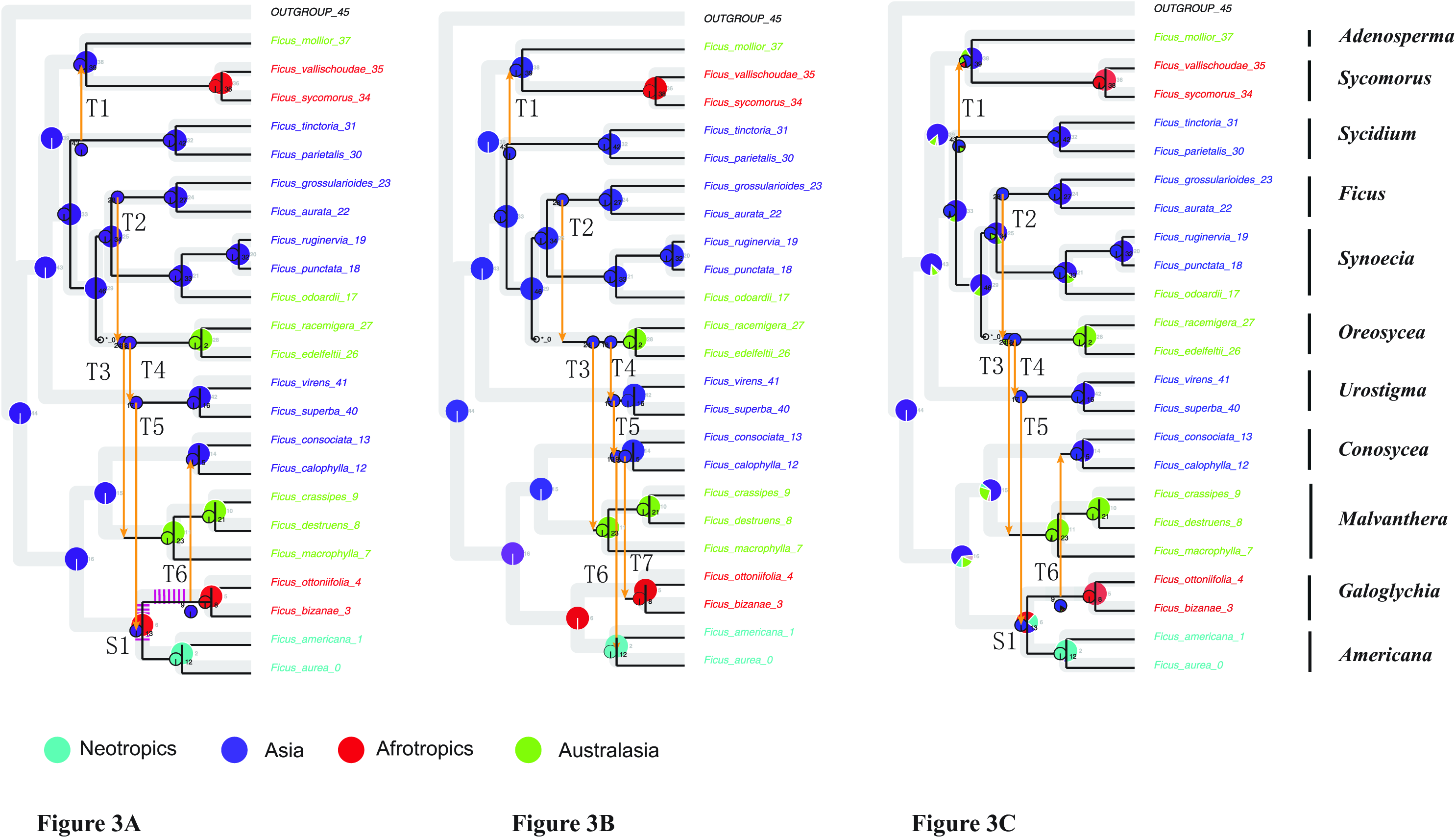
Results of the reconciliations inferred by *Mowgli* on the fig/fig wasp data set with default cost settings: A) using a single most likely area for ancestral species and without enforcing geographic constraints (events: 17 S, 6 T, 1 L), purple dashed branches indicate parts of the reconciliation where geographic constraints are not fulfilled; B) using a single most likely area when enforcing geographic constraints (events: 16 S, 7 T, 1 L).The leaves of the phylogenetic trees are coloured according to current geographic distribution of the associates. In both cases, big pie charts correspond to the *Ficus* ancestral geographic areas, small pie charts correspond to the pollinator ancestral geographic areas; C) Reconciliation obtained when alternative ancestral areas are considered (namely those with probability>0.15). Sections of the pies are proportional to the probability of the associated states. *Ficus* taxonomic subdivisions are reported on the right inside of the figure.

## 4 DISCUSSION

We provide here significant extensions for a reconciliation tool (*Mowgli*) and a visualization tool (*Sylvx*) to infer co-diversification scenarios that, for the first time, can take the historical biogeographies of the associated lineages into account. The extension of the *Mowgli* software precludes geographic inconsistency during the reconciliation process. The resulting reconciliations can then be visualized and edited in the *SylvX* updated graphical interface that now integrates annotations of ancestral nodes. *Mowgli* is already one of the few time-consistent efficient methods that build optimal reconciliations. With the integration of geographic constraints in its algorithm, this tool now provides more realistic codiversification scenarios than other reconciliation methods. Producing biologically realistic scenarios can ease their interpretation. In addition, geography-aware reconciliations can reveal whether host switches occur in sympatry or whether they are associated with dispersal events of the symbionts/parasites: this helps unravelling the evolutionary processes underlying host switches

In the particular example of the fig/fig wasp interaction presented here, the geographic inconsistency revealed at one of the cospeciating nodes in the analysis (Fig 3A) ran without constraints might actually point out some ambiguity in the biogeographic history of the *Ficus* hosts. According to the inference conducted in Cruaud *et al.* (2012) the most likely area for the common ancestor of Neotropical figs (belonging to the *Americana* section) and Afrotropical figs (belonging to the *Galoglychia* section) is Africa while the proposed cospeciating pollinators lived in Asia (S1; Fig. 3A). In order to respect geographic consistency (when only the most likely area is kept for each ancestral species, Fig. 3B), our geography-aware reconciliation suggests that the current association of figs wasps with *Galoglychia* in Africa, resp. *Americana* in the Neotropics, is the result of two independent switches (Fig. 3B, events T6 and T7) of the pollinators from an Asian fig ancestor (the ancestor of the *Conosycea* figs). However, the biogeographic analysis of Cruaud *et al.* (2012) also suggested that the node S1 of *Ficus* could be situated in Asia (though with a much lower likelihood than the Afrotropics). When specifying alternative geographic areas (Fig. 3C), including Asia for the conflicting node in the *Ficus* phylogeny, we obtain a reconciliation that matches the one obtained without constraints (therefore entailing one less transfer and one more cospeciation event). This result suggests that the common ancestor of the African figs of section *Galoglychia* and the new world figs from the section *Americana* could have been located in Asia. Under this latter scenario most of the host switches observed happen in sympatric settings. We will not conclude on the biogeographic history of the fig/fig wasp association as the purpose of our study is not to explore alternative scenarios for this association. The above discussion mainly demonstrates the utility of our method in revealing inconsistency between biogeographic scenarios and a cospeciation hypothesis and therefore proposing alternative scenarios that conciliate both. As in all ancestral character state inferences that rely on present day data, biogeographic reconstructions entail some incertitude. In particular, they are highly sensitive to missing data (species that have not been sampled and/or extinct species). It is therefore important to compute reconciliations with alternative ancestral ranges to investigate biogeographic scenarios.

### Perspectives

The tools developed in this study can be applied to all interspecific interactions for which biogeographic scenarios are available for both partners. Fast developments in sequencing technologies generate more accurate and more exhaustive phylogenies and methods in historical biogeography have also improved. Therefore, we can hope that numerous datasets will be available in the near future and that cospeciation could be tested on more systems (Cruaud & Rasplus 2016). For instance, robust phylogenies and biogeographic scenarios are now available for groups of lice that have been model systems in coevolutionary studies (Boyd *et al.* 2017). Once comprehensive phylogenies of the hosts are available, our method could be used to better understand the geographic context of host switches in this model system. Geography-aware reconciliation could also be applied to explore the diversification history of the numerous parasitic wasps that are part of the microfauna exploiting figs: several lineages of parasitic wasps have been shown to partly cospeciate with their host figs (Jousselin *et al.* 2008; Jousselin *et al.* 2006) and biogeographic scenarios for some lineages are available (Cruaud *et al.* 2011). These developments could also be applied to specific sections of the genus *Ficus* in order to shed light on their complex biogeographic histories (e.g. section *Urostigma* that has experienced several dispersal events between Africa and Asia, Chantarasuwan *et al.* 2016). Other nursery pollination/mutualisms such as the interaction between *Yucca* and their pollinating moths are also good candidates for including geographic constraints into coevolutionary scenarios, as some studies have questioned the respective role of geography and host-plant association in driving the diversification of *Yucca* moths (Althoff *et al.* 2012). Plant/pollinator systems (Hutchinson *et al.* 2017), parasitoid/host insect associations (Deng *et al.* 2013; Wilson *et al.* 2012), herbivorous insect/plant interactions (*e.g.* McLeish *et al.* 2007; Percy *et al.* 2004) and various vertebrate/parasite associations (*e.g.* Badets *et al.* 2011; Bentz *et al.* 2006; Weckstein 2004) for which researchers have investigated the relative role of geography and biotic interactions in shaping cophylogenetic signals could also be studied.

Furthermore, the approach presented in this paper does not only apply to geographic information and could be extended to other biological traits. For instance in systems where the species are partitioned into different habitats (*e.g.* forest canopy species vs savannah species), geographic areas could be replaced by traits related to the ecological niches; constraints that are similar to the ones applied for geography could then be easily transferrable. Informing ancestral characters for habitats on the host and the symbiont phylogenetic trees and using “*Mowgli* with constraints” would result in constraining cospeciation and host switches to associates sharing the same ecological habitats. In a similar way, the respective climatic niches of associated organisms could also be used when parasite (or symbiont) distributions are known to be strongly constrained by thermal tolerance (see Singh *et al.* 2017, for a recent study showing that climatic conditions influences the patterns of association between fungi and their algal partners). In many specialized interactions, such as host/obligate bacterial endosymbionts (*e.g.* Jousselin *et al.* 2009, Rosenblueth *et al.* 2012) or host/viruses associations (Ramsden *et al.* 2009; Garamszegi 2009), inferring ancestral character states for some ecological traits for the “symbiotic” lineages (the parasite) independently of their hosts is not always straightforward. However, the evolution of these obligate associations and their maintenance are still governed by some phenotype matching between the partners. For instance in host/bacterial symbiont associations, the metabolic complementarity of the host and the symbiont (Zientz *et al.* 2004) could be reconstructed and used to constrain the reconciliations. In host/virus associations, information about the host immune system and viruses adaptations could be used (Longdon *et al.* 2014). The extension of *Mowgli* proposed here could probably be adapted to fit the biological properties of these associations

Independently of the new functionality implemented in *Mowgli*, the concomitant update of *SylvX* allows the comparison of ancestral states for any character of the hosts and/or the symbionts. This can help interpreting reconciliations by replacing them in their biological context. One of the most useful functionalities of *SylvX* is now to be able to visualize whether host switches are associated with evolutionary transitions in character states in both the parasite and/or the host. It can therefore help understanding the biological processes that are associated with these transfers. Mapping characters of the associates throughout the reconciliation can also help investigating whether there is correlated trait evolution in host and parasites. Until now, such correlations could only be investigated on one of the associate phylogeny (*e.g.* Sorci *et al.* 2003; Jousselin *et al.* 2003). Looking at simultaneous transitions in character states in both partners throughout a host/parasite reconciliation might help identifying co-adapted traits that constrain the association.

In conclusion, we provide here a framework that can integrate the character histories of the associates into the reconciliation process. It can take into account incertitude in the character states and allows recovering biologically realistic scenarios. It can also shed light on character history inferences by pointing out inconsistencies between the character states of the two associates on the reconciliation map. The new developments made in *SylvX* facilitate these interpretations. A more integrative approach than the one presented here, would consist in co-optimizing the reconciliation and the biogeographical inference simultaneously. However this would require using the same optimization criterion for both inferences and setting adequate parameters for these very different processes in a single model. When conducted, this work should probably rely on Maximum Likelihood optimization as in the ALE reconciliation software (Szöllősi *et al.* 2012). For now, we believe that the use of “constraint-aware” reconciliations is preferable to current practices that consist in elaborating ad-hoc narratives once the reconciliations are obtained and compared with the character histories of the associates.

## ACKNOWLEDGMENTS

*Funding*: ANR Phylospace. We thank A. Cruaud, JY Rasplus & coll. for sharing the fig/fig-wasp dataset.

## AUTHOR CONTRIBUTION

VB, JPD and EJ designed the study. VB and JPD developed *Mowgli*, FC developed *SylvX*. VB and EJ wrote the manuscript with contributions of FC.

## SOFTWARE AVAILABILITY

Mowgli is available on http://www.atgc-montpellier.fr/Mowgli/ with manual, *GeoRecHelper* and example files. It runs on OSX (Mac) and Linux systems.

*SylvX* is available on www.sylvx.org with manual and example files and can be installed on any platforms.

Supplementary Material 1: Description of the pipeline to generate trees and annotation files that can be taken as inputs for both *Mowgli* and *Sylvx*.

Supplementary Material 2: Reconciliations obtained under different cost settings on the mock dataset.

